# Innovations to expand drone data collection and analysis for rangeland monitoring

**DOI:** 10.1101/2021.02.05.430004

**Authors:** Jeffrey K. Gillan, Guillermo E. Ponce-Campos, Tyson L. Swetnam, Alessandra Gorlier, Philip Heilman, Mitchel P. McClaran

## Abstract

In adaptive management of rangelands, monitoring is the vital link that connects management actions with on-the-ground changes. Traditional field monitoring methods can provide detailed information for assessing the health of rangelands, but cost often limits monitoring locations to a few key areas or random plots. Remotely sensed imagery, and drone-based imagery in particular, can observe larger areas than field methods while retaining high enough spatial resolution to estimate many rangeland indicators of interest. However, the geographic extent of drone imagery products is often limited to a few hectares (for resolution ≤ 1 cm) due to image collection and processing constraints. Overcoming these limitations would allow for more extensive observations and more frequent monitoring. We developed a workflow to increase the extent and speed of acquiring, processing, and analyzing drone imagery for repeated monitoring of two common indicators of interest to rangeland managers: vegetation cover and vegetation heights. By incorporating a suite of existing technologies in drones (real-time kinematic GPS), data processing (automation with Python scripts, high performance computing), and cloud-based analysis (Google Earth Engine), we greatly increased the efficiency of collecting, analyzing, and interpreting high volumes of drone imagery for rangeland monitoring. End-to-end, our workflow took 30 days, while a workflow without these innovations was estimated to require 141 days to complete. The technology around drones and image analysis is rapidly advancing which is making high volume workflows easier to implement. Larger quantities of monitoring data will significantly improve our understanding of the impact management actions have on land processes and ecosystem traits.

## Introduction

The rangeland manager’s challenge is the extensive management across a heterogeneous landscape under an uncertain climate. With so much uncertainty, rangeland managers typically opt for an adaptive management approach, particularly in the public domain rangelands that dominate the western US. Adaptive management is not simply trial and error, but according to the Department of Interior (Williams et al., 2009): *An adaptive approach involves exploring alternative ways to meet management objectives, predicting the outcomes of alternatives based on the current state of knowledge, implementing one or more of these alternatives, monitoring to learn about the impacts of management actions, and then using the results to update knowledge and adjust management actions*. Unfortunately, budgetary and institutional constraints have long limited public land monitoring, as noted by Fernandez-Gimenez et al. (2005). Sayre et al. (2013) state that *monitoring is a critical component of adaptive management but often weak or missing in practice*. The premise of this paper is that expanded monitoring is a prerequisite for improved rangeland management.

Traditional field monitoring methods (e.g., transects or quadrats) can provide detailed information for assessing the health of rangelands. Cost, however, often limits monitoring locations to a few key areas or random plots that observe a small fraction of the land they are intended to represent (Booth and Cox, 2011; Toevs et al., 2011; West, 2003). Remotely sensed imagery enables a broader view of the land and potentially a more representative sample. Drone-based imagery, in particular, can observe larger areas than field methods while retaining high enough spatial resolution to estimate many rangeland indicators of interest. These indicators include vegetation cover (Baena et al., 2017; Breckenridge et al., 2011; Hardin et al., 2007; Laliberte and Rango, 2011), vegetation heights (Cunliffe et al., 2016; Gillan et al., 2020; Jensen and Mathews, 2016; Olsoy et al., 2018), biomass (Cunliffe et al., 2016; Michez et al., 2019), forage utilization (Gillan et al., 2019), and soil erosion (D’Oleire-Oltmanns et al., 2012; Gillan et al., 2017).

At present, leveraging small drones, off-the-shelf sensors, and structure-from-motion photogrammetry (SfM-MVS) is a low-cost workflow capable of meeting several rangeland monitoring needs. However, challenges remain to deploy this technology at larger operational scales. The geographic extent of drone imagery products is often limited to a few hectares (for spatial resolution ≤ 1 cm) due to image collection and processing constraints. Additionally, sharing data and reporting out monitoring results to collaborators and stakeholders can be limited by large file sizes and the complexity of web development. Overcoming these limitations would move us closer to realizing the potential value of drone-based monitoring, which is: 1. broader extent observations; 2. better measurement of some indicators; and 3. permanent visual records. Scaling the production and interpretation of drone imagery will be essential to support adaptive management on individual allotments as well as to integrate with national-scale monitoring programs such as the Bureau of Land Management’s Assessment, Inventory, and Monitoring (AIM) strategy and the Natural Resource Conservation Service’s National Resource Inventory (NRI).

Our objective was to develop a workflow to increase the extent and speed of acquiring, processing, and analyzing drone imagery for repeated monitoring of two common rangeland indicators: vegetation cover and vegetation heights. We compared the total number of workdays to execute our innovative workflow with the time required to complete a more conventional workflow. We then demonstrate sharing and visualization of the imagery products and results using free or open-source web tools. We focused on the workflow and did not directly assess the accuracy of indicator values compared with field methods. The workflow described here is an initial phase of a larger research project investigating the use of drone imagery for mapping ecological states (Steele et al., 2012).

## Methods

### Study Area

We conducted this research at Santa Rita Experimental Range (SRER) in southern Arizona (31°48’36”N, 110°50’51”W; Fig. 1). The range, established in 1902, is a 21,000 ha Sonoran Desert grassland that has been significantly invaded by velvet mesquite (*Prosopis velutina*). SRER is a living laboratory for studying dryland ecology and sustainable livestock production. The range has over 200 permanent long-term transects intended to capture vegetation dynamics across multi-decadal time spans (McClaran et al., 2002; cals.arizona.edu/srer). In the upper elevations of the range (1050-1300 m MSL; Major Land Resource Area 41-3), we selected a subset of 100 transects for this study. The long-term transect locations are not randomized and thus do not represent an unbiased sample of the study area. It was not our intent to extrapolate results to monitor all of SRER. Instead, the legacy transect locations provided a large sample size from which to demonstrate our workflow.

**Fig. 1.**
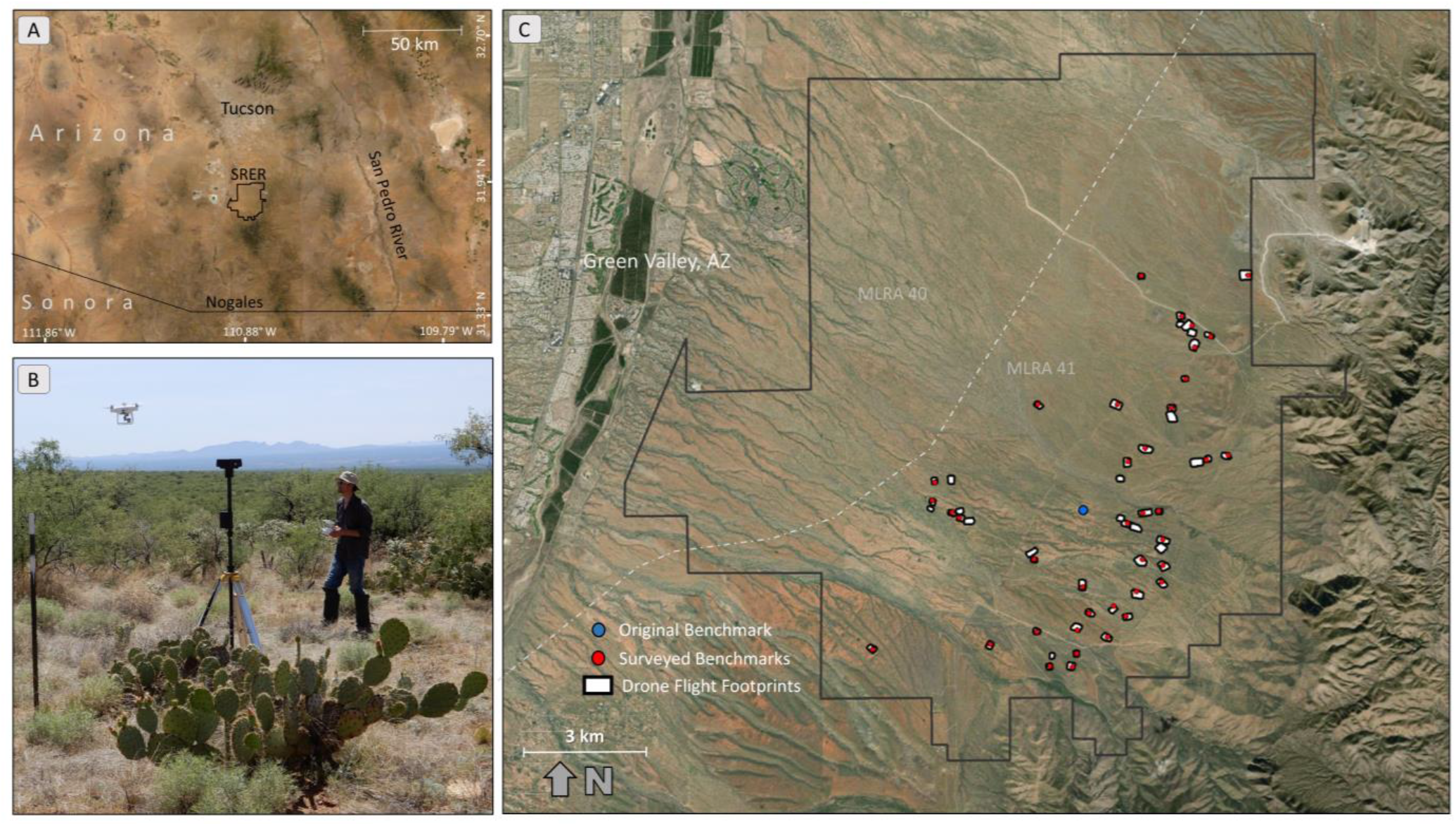
A) This project occurred at Santa Rita Experimental Range (SRER) in southern Arizona. B) We collected aerial imagery using a DJI Phantom 4 RTK with portable base station. C) We collected imagery at 53 flight plots covering a total of 193 ha in May 2019 and repeated in September 2019. The drone was launched near surveyed benchmarks (shown as red points).

### Image Acquisition

We collected drone imagery covering the 100 transects in May 2019 (dry season) and repeated the acquisition in September 2019 (monsoon season). We used a DJI Phantom 4 RTK quadcopter specifically because it possessed a real-time kinematic global navigation satellite system (**RTK GNSS**). RTK GNSS on drones is not a new technology, but it is now more accessible due to its integration in off-the-shelf aircraft at reduced cost. The Phantom 4 RTK in 2019 cost ~$8,000 and came paired with a portable GNSS base station and tripod (D-RTK 2).

RTK is a technology that pinpoints the 3D coordinates of the camera for each image taken from the moving drone. It can be accurate within a few centimeters, which is more precise than a typical global positioning system (GPS) receiver is. RTK GNSS is a differential correction system where the aircraft is in constant communication with a nearby portable base station with known coordinates (i.e., placed over a surveyed benchmark). When an image is taken, the location of the drone (and more specifically the camera), as estimated from the onboard GPS, is compared with and corrected by a signal from the base station. The improved location coordinates (i.e., latitude, longitude, elevation) are then recorded as metadata on the exchangeable image format (EXIF) header of each image.

Highly accurate camera locations can replace the use of ground control points (GCPs) to scale and georeference imagery products such as point clouds and orthomosaics (Forlani et al., 2018; Hugenholtz et al., 2016; Rehak et al., 2013). RTK allowed us to streamline two aspects of the workflow. First, it eliminated the need to place and survey GCPs with either a total station or ground-based differential GPS. It can be quite cumbersome to survey GCPs, especially for large flight areas that may require a dozen or more. Second, labor was eliminated in the photogrammetry processing step of identifying each GCP in every image. Algorithms in commercial software aimed at automatically identifying GCPs are not always successful, especially for oblique angle views. With RTK drones, we can collect and create high-quality image products over large extents, while a GCP workflow practically limits us to plot scales.

Prior to this study, SRER had only one known surveyed benchmark. We established and surveyed more benchmarks using a Trimble R10 RTK GNSS (base station and rover). We set the Trimble base station over the original benchmark and roved across the range setting up new benchmark points near all of the flight transects. Because of some transect clustering, we needed just 39 benchmarks to cover the 100 transects (Fig. 1). The benchmark points were existing rebar posts that marked the ends of long-term transects. Absolute accuracy of the surveyed benchmarks was < 1 cm horizontal and 1-1.5 cm vertical. We used the drone portable base station (D-RTK 2) placed over the benchmarks to facilitate RTK location correction while the drone flew and collected images.

Through our own independent assessment, we found the RTK drone imagery products (flown at 38 m above ground level) to have horizontal location accuracy of 2.2 cm and vertical accuracy of 3.4 cm. This was within ~1 cm, both horizontally and vertically, of an assessment conducted by DroneDeploy (Mulakala, 2019). Our reproducibility assessment yielded a horizontal precision of 3 cm and vertical precision of < 1 cm for digital surface models.

For each of the two campaigns (dry and wet seasons), we collected 53 flight plots to cover the 100 transects, a total of 193.1 ha (Fig. 1). Transects that were very near each other (< 300 m) were often captured in a single image product. Flight plots ranged in size from 1.6 to 7.1 ha to meet the objectives of the ecological state mapping project. We collected a high density of nadir and oblique images (~200 ha^-1^) in order to create very detailed and accurate point cloud models and downstream products such as vegetation height models (VHMs). See Table 1 for full sensor and acquisition specifications and Fig. 2 for a chart of the entire workflow.

**Table 1.**
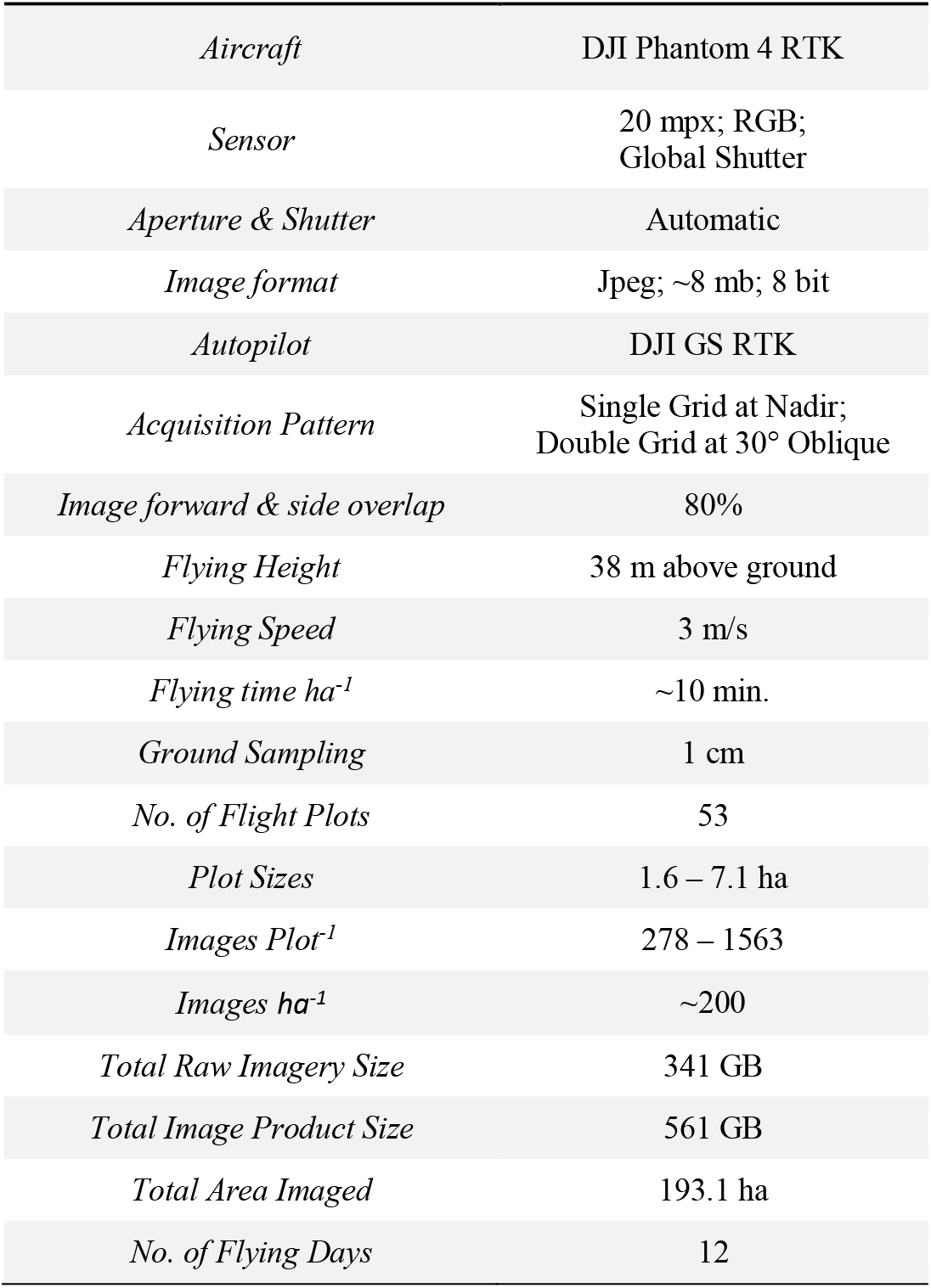
Hardware and image acquisition specifications for the data collection campaigns that occurred in May 2019 and repeated in September 2019

**Fig. 2.**
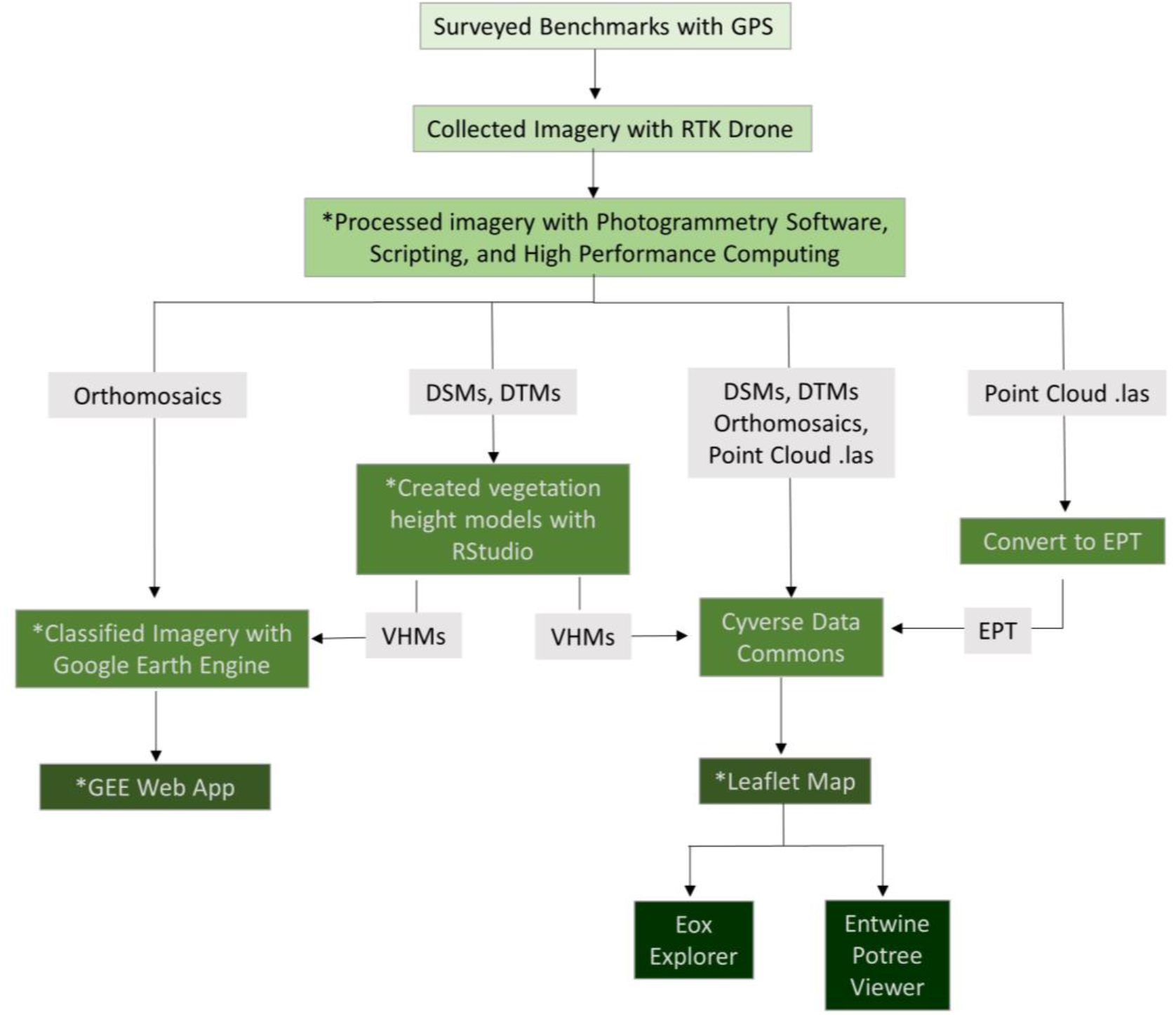
Workflow for data collection, processing, and sharing. DSMs = digital surface models; DTMs = digital terrain models; VHMs = vegetation height models; EPT = entwine point tile. Items with * have available code.

### Image Product Creation

Eliminating ground control points through the use of RTK enabled us to **fully automate imagery product creation with Python scripts**. What would take an analyst a few hours to complete interactively (in addition to the dense point cloud reconstruction time), was scripted in Agisoft Metashape 1.5.2 (www.agisoft.ru). The general SfM-MVS workflow is well documented so it will be abbreviated here (see Eltner et al., 2015; Smith et al., 2015; Snavely et al., 2008; Westoby et al., 2012). Python scripts, running from command line, added imagery to the project, created the sparse point cloud, filtered poor quality points, optimized the sparse model, then generated dense point clouds, digital surface models, digital terrain models, and orthomosaics (see Table 2 for processing parameters). When the plot completed, it seamlessly started the next plot. Image processing reports were later spot checked for quality assurance.

**Table 2.**
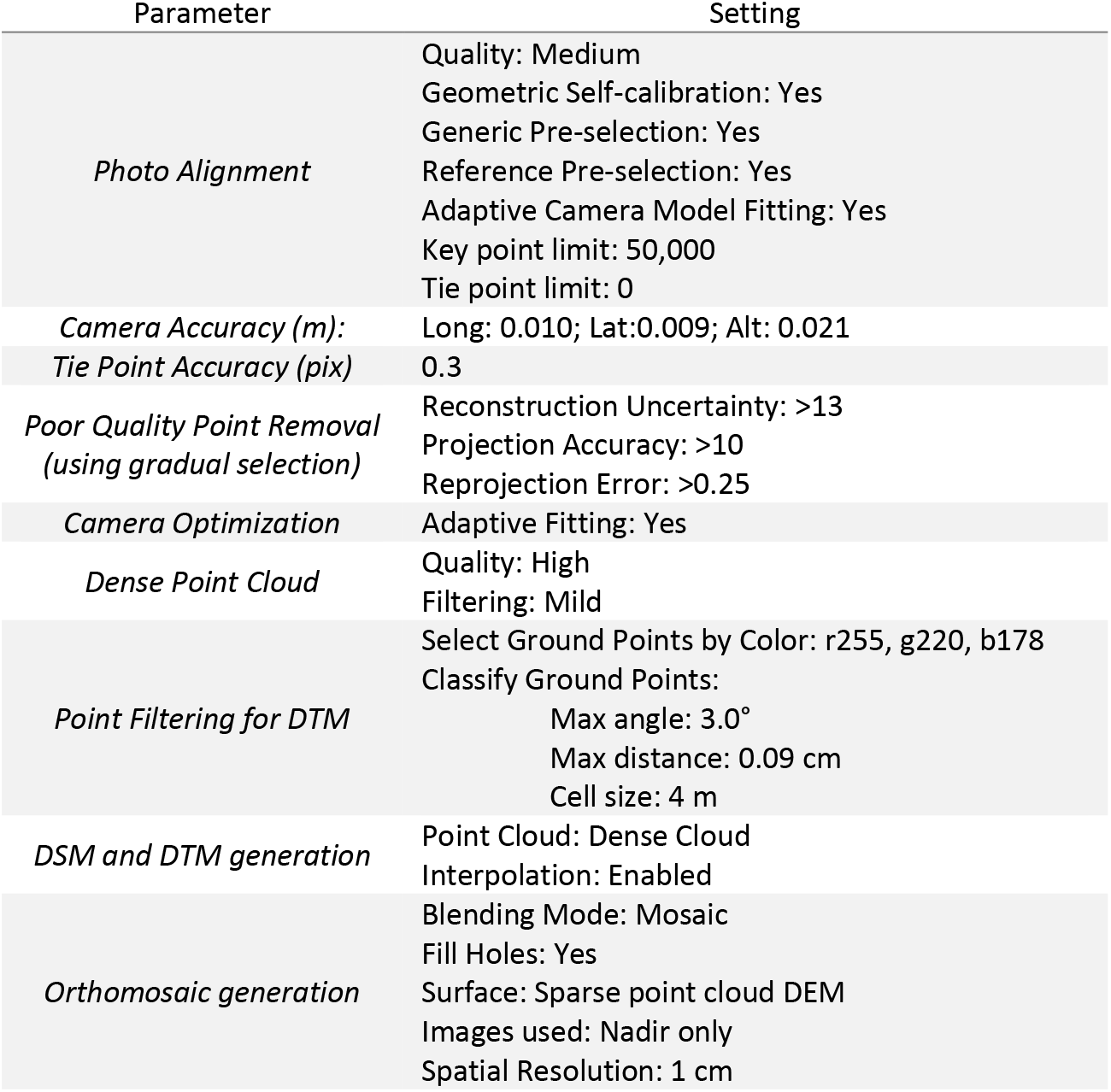
Structure-from-motion photogrammetry processing parameter settings using Agisoft Metashape 1.5.2.

In addition to scripting, we used **high performance computing (HPC)** to quicken image product creation for the twenty largest flight plots (801-1600 images each). We used the University of Arizona HPC system called Ocelote. Each CPU node was an Intel Haswell V3 28 core processor with 192 GB RAM. They also had Graphical Processing Units (GPU) nodes with one Nvidia P100 GPU, 28 cores, and 256 GB RAM. The type and number of nodes used depended on the availability of HPC resources. We typically used between 10-15 nodes working in parallel, each running an instance of Metashape, which was designed with network processing in mind. Each Metashape instance and license operated through container software Singularity (singularity.lbl.gov). Containers enabled us to package our computing environment, including software installs and licenses, for easy deployment on the remote HPC nodes. We had to purchase educational Metashape licenses for each processing node (~$500 each). Our Metashape instance ‘master’ was located on a Linux server while the worker nodes were provided by the HPC (also Linux). For the 33 smaller plot areas (278-800 images), we used a Windows desktop machine (hereafter as the PC) with two Intel Xeon CPUs (2.4 GHz; 16 logical processors each), two Nvidia GeForce GTX 1080 video cards (GPUs), and 256 GB RAM.

Using both the HPC and the PC simultaneously, it took approximately two weeks to produce the entire suite of imagery products (point clouds .las, digital terrain models .tif, digital surface models .tif, orthomosaics .tif) for one collection campaign, a total of 561 GB (Fig. 3). We then generated vegetation height models (VHMs) for each plot area by subtracting the digital terrain model from the digital surface model on a cell-by-cell basis using the *Raster* package in Rstudio. This was executed on the PC and took approximately 4 hours to complete. With a simple shell command (see https://entwine.io/quickstart.html), we converted all of the .las point clouds to entwine point tile (EPT), a format that facilitates browser-based viewing of large point clouds. We uploaded all image products and raw imagery to Cyverse Data Commons (cyverse.org/data-commons/....pendingDOI) for public access and long-term storage.

**Fig. 3.**
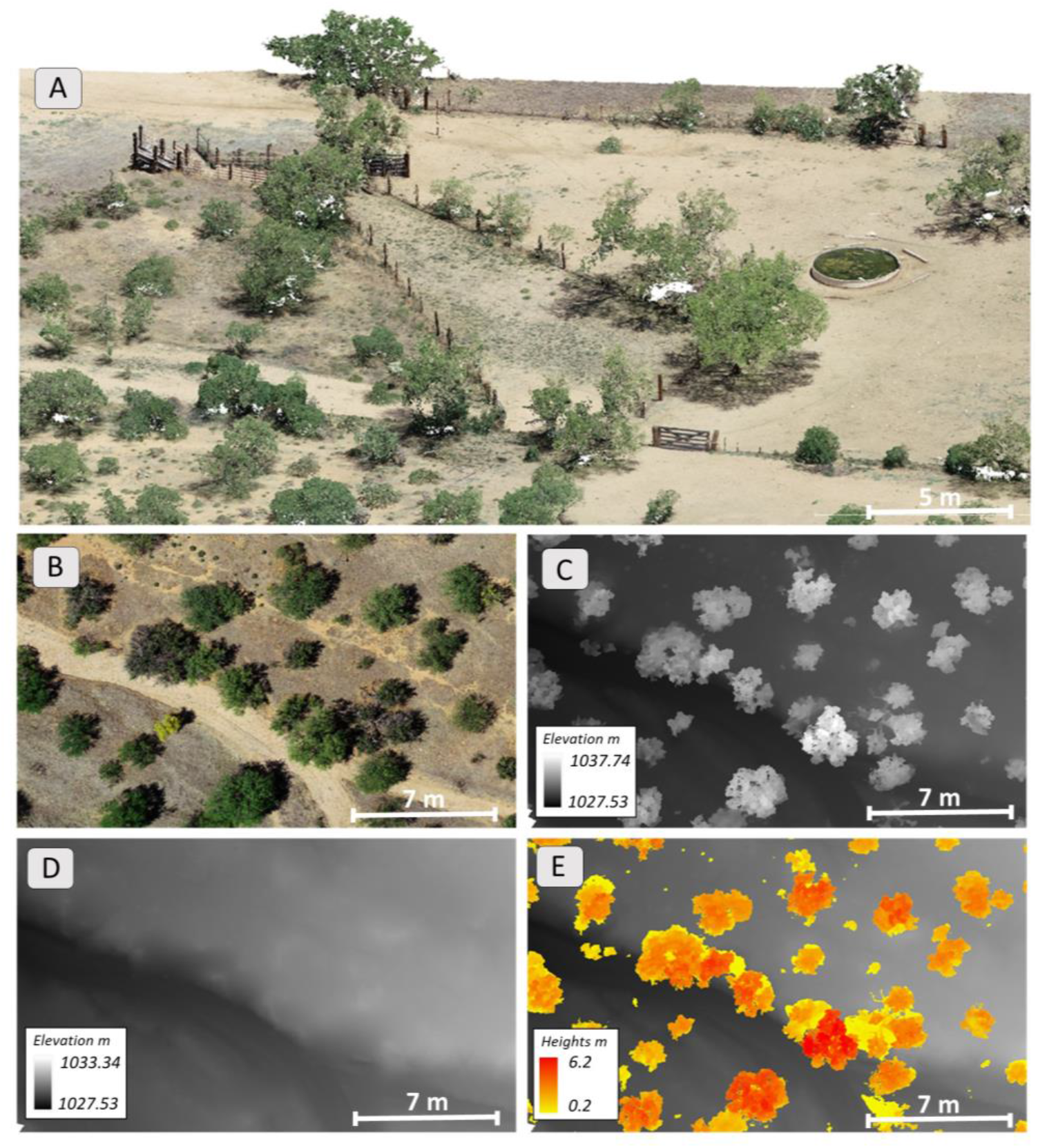
Imagery products created from drone imagery, including A) Dense point cloud; B) True-color Orthomosaic; C) Digital surface model; D) Digital terrain model; and E) Vegetation height model.

### Image Product Analysis

As large drone imagery datasets outpace desktop computing power, new tools are needed for rapid analysis, visualization, and sharing. We used the **cloud-based analysis platform Google Earth Engine (GEE; earthengine.google.com**) to derive additional value-added indicators from the imagery products. GEE is a cloud-based geospatial analytics platform with access to large computational resources and two application programming interfaces (API), JavaScript and Python. These APIs provide a suite of raster analysis functions including several classification algorithms (Gorelick et al., 2017). Though it was built primarily for broad scale satellite imagery, it is free and can also handle very large drone datasets. A powerful feature of GEE is the ability to easily share JavaScript code and imagery assets between users, which can make imagery analysis collaborative.

We uploaded all orthomosaics from the May acquisition (n=53) into GEE and then mosaicked them together to form a single large super-mosaic (19.3 billion pixels). We repeated these steps for the May VHMs, September orthomosaics, and September VHMs. We used red, green, and blue bands, vegetation heights, and a calculated green leaf algorithm (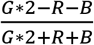; Louhaichi et al. 2001) as input features to thematically classify the imagery with a machine learning classification tree algorithm (Breiman et al., 1984). We identified four cover classes as a simple demonstration of the tool and workflow: herbaceous vegetation, woody vegetation (including cactus), bare-ground, and shadow. We used the polygon digitizing tool within GEE to select training data for each class. We generated seven training polygons for each class, with each training polygon containing hundreds of training pixels. For classification validation, we randomly selected 50 pixels for each class across the super-mosaic. These pixels were visually interpreted and compared with their assigned class.

For comparison with a conventional workflow, we classified the drone imagery using ArcGIS Pro 2.5 (esri.com) installed on the PC. We used the same input features and basic training procedures as our GEE workflow. Instead of merging all the orthomosaics into a super-mosaic (as we did in GEE), we used *Model Builder* to automate the sequentially classification of each orthomosaic using the *Random Trees* algorithm. We enabled parallel processing to use all available CPUs for faster classification.

### Visualization and Sharing

For sharing monitoring results and image product visualization on the web, we chose two platforms. We developed a public facing web-app directly in GEE that enables users to view the orthomosaics, VHMs, classified maps, and see summaries of the vegetation cover and vegetation heights. The website was developed with JavaScript and is served through Google Cloud. Additionally, we developed a mapping application using Leaflet, an open-source JavaScript library (https://de.cyverse.org/....pendingDOI). Users are able to explore a map of all the flight plots at SRER. Clicking on individual plots invites users to view high-resolution versions of the orthomosaics and 3D point clouds directly in their web browser. The orthomosaics are displayed in Eox Cog Explorer (https://geotiffjs.github.io). The point clouds are viewable using Potree (entwine.potree.io), a free open-source web graphics library that renders point clouds directly in your web browser using the EPT format.

## Results and Discussion

By incorporating a suite of existing technologies in drones (RTK GNSS), data processing (automation with Python scripts, high performance computing), and cloud-based analysis (Google Earth Engine), we increased the efficiency of collecting, analyzing, and interpreting high volumes of drone imagery for rangeland monitoring. End-to-end, our workflow took 30 days, while a workflow without these innovations was estimated to require 141 days to complete (Table 3).

**Table 3.**
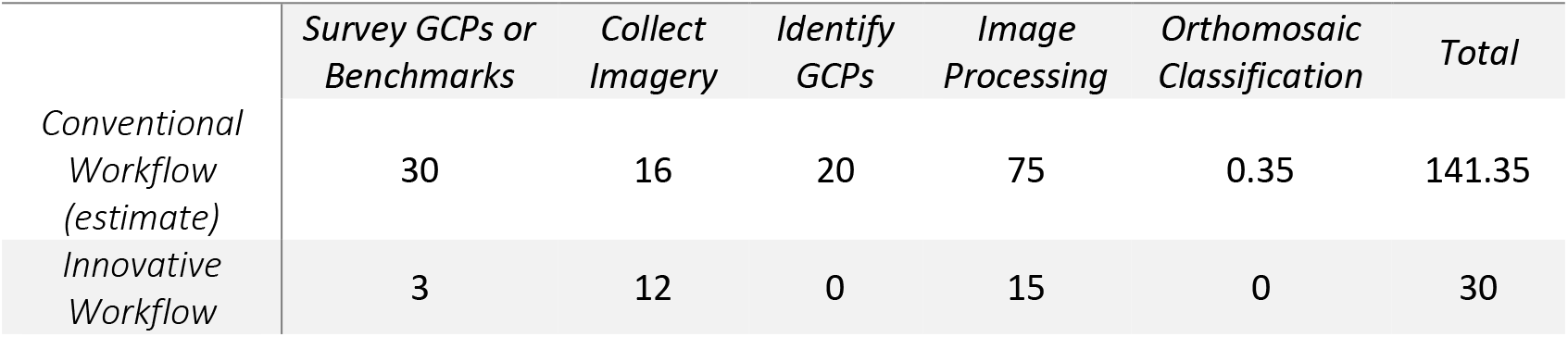
Number of workdays to collect, process, and analyze drone imagery collected in May 2019

RTK saved us considerable time in the image collection step (Table 3). With a GCP workflow, small plots would require 5 to 8 GCPs, and larger plots could require 10 to 20 GCPs to achieve accuracies comparable to the RTK results (James et al., 2017; Sanz-Ablanedo et al., 2018). A conservative estimate would be 300 GCPs for all of the flight plots, which could take upwards of 30 workdays to install and survey. Our RTK workflow, for comparison, required just 3 days to survey 39 benchmarks at existing stakes. Placing and collecting GCP targets before and after the flights would add an additional ~30 minutes to each plot. This could push the total number of flying days from 12 (with RTK) to 16. Our RTK workflow eliminated the manual labor of identifying GCPs during the image processing step, which could take hours per plot. We estimated a savings of 20 workdays by eliminating manual GCP identification.

Other potentially more efficient options for image product referencing exist. For example, cellular tower virtual reference systems can send correction signals to flying drones using tablets or smartphone devices as an intermediary. These correction networks could eliminate our need to use portable base stations and surveyed benchmarks. In Southern Arizona, a private company provides the correction signal as a service, but we decided against this option because strong cellular reception was not reliable everywhere in the study area. As cellular coverage expands, even across rural rangelands, virtual reference systems will become increasingly viable for drone image product referencing. Alternatively, the drone and portable base station workflow used in this project could be executed without surveying benchmarks. In remote areas where high-precision surveying is not practical or the equipment is not available, drone image products can be corrected to have high *relative* accuracy. In this case, the image products are correctly scaled but may be shifted horizontally or vertically from a true absolute position (see Gillan et al., 2020).

The HPC was 14 to 24x faster than the PC at dense point cloud reconstruction, depending on the number of HPC nodes and the total number of images in the Metashape project. Plots with larger numbers of images required much greater (non-linear increases) processing time and showed the most speed gains through the HPC. For example, a plot with 900 images that took 24 hours to process on the PC, was completed in 1.6 hours on the HPC. A 1500 image plot that took 120 hours to process on the PC, was completed in 5 hours on the HPC. By using the HPC on the twenty largest plots, we saved ~45 days of image processing. Additionally, scripting increased the speed of processing the plots on the PC by processing 24 hours per day including starting jobs in the middle of the night. This probably saved ~15 days.

In the near future, computational power will not be a hindrance to high volume drone data. For example, recent software updates to Agisoft Metashape (v. 1.6.2) have significantly increased the speed of image processing on PC and HPCs. We can now expect the processing time to be 3-8x faster than described in this paper. HPC is becoming increasingly available through many universities with easier to use interfaces (Settlage et al., 2019). Alternatively, image processing can be outsourced (via the web) to commercial entities including DroneDeploy (dronedeploy.com), Pix4D (pix4d.com), and Delair (delair.aero).

Classifying all drone orthomosaics in GEE was essentially instant. Near instant feedback allowed us to quickly assess classification results and adjust training data for higher accuracy (see Appendix Tables S1 & S2 for confusion matrices). In comparison, it took ~3 hours to classify 53 orthomosaics using ArcGIS Pro on the PC.

GEE worked well for classifying the imagery and is currently the most mature tool for quickly analyzing large quantities of drone imagery. However, limitations of the platform include data storage limits and upload/download speeds to and from GEE. Additionally, it has limited functionality to conduct every analysis we might want for rangeland monitoring (e.g., 3D point cloud analysis; landscape metrics). A greater variety of analysis options exist in ArcGIS Pro, but they may be less accessible to users due to cost. Fortunately, there is an enormous and growing variety of image analysis tools available across open platforms such as R, Python, and QGIS. Many have the capability to maximize local computing resources and distribute processing tasks to HPC clusters (see parallel processing options for R and Python). The availability of high throughput analysis tools will soon not be a constraint. Instead, the challenge will be to identify workflow ‘best practices’ for estimating a suite of rangeland indicators and selecting the best mix of tools that are cost-effective and repeatable (Gillan et al., 2020).

Leaflet paired with Eox COG Explorer and Potree provided an easy-to-build web map for visualizing the point cloud and orthomosaic products (Fig. 4; https://de.cyverse.org/....pendingDOI). The Potree viewer has basic analysis tools (distance, volume, profile). The GEE app enabled us to share the classified maps, VHMs, and graphed summaries of vegetation cover and heights (Fig. 5; https://bit.ly/srer-drone-2019). Both of these sharing options eliminated the need for collaborators to download large files or install 3^rd^ party software on their local machines.

**Fig. 4.**
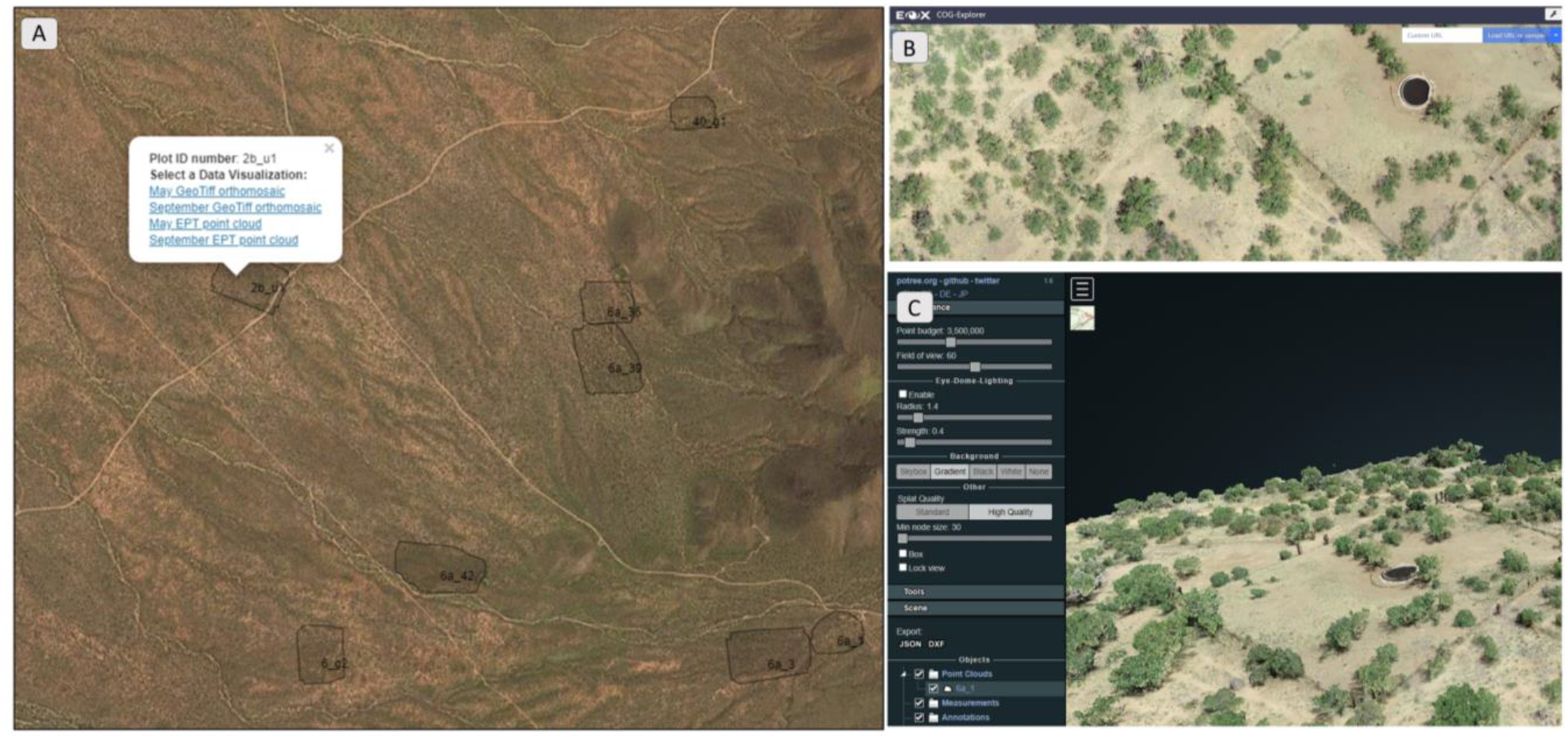
A) We created an open-source Leaflet map to enable collaborators to view imagery products through a web-browser (https://de.cyverse.org/....pendingDOI).B) High-resolution orthomosiacs can be viewed with Eox COG Explorer. C) Point clouds can be viewed with a Potree viewer.

**Fig. 5.**
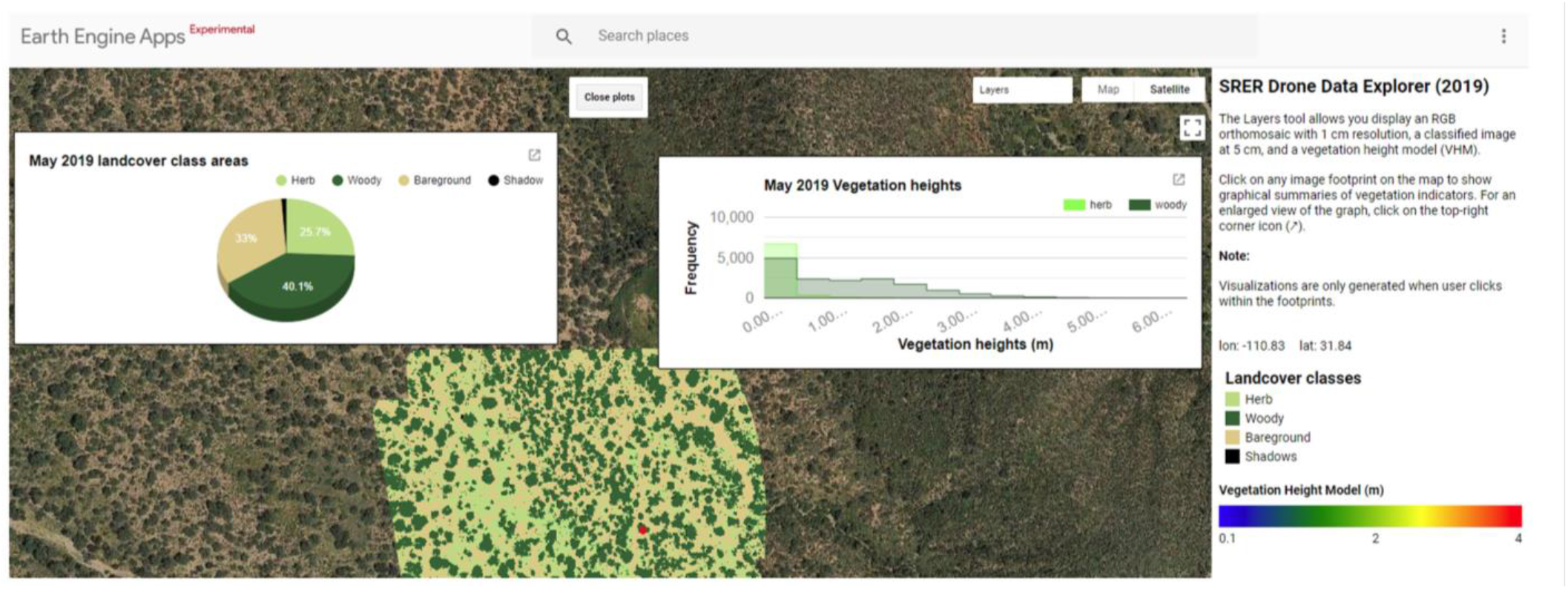
We developed Google Earth Engine web-app showing classified maps, vegetation height models, and indicator summaries for vegetation cover and heights. https://bit.ly/srer-drone-2019

### Implications

High volume drone imagery will enable us to move beyond ‘proofs of concept’ and other small-scale research demonstrations to data quantities that significantly improve our understanding of land processes. In an adaptive management framework, this means expanding monitoring beyond the confines of plots and transects to provide a more representative sample of vegetation characteristics across rangelands. A more representative sample could increase the statistical power to detect indicator change by either increasing the sample size (i.e., collecting imagery at more locations than transects), or by expanding the observational area of each transect to reduce variance between samples (Sundt, 2002). Our drone imagery covered 193 ha during the dry and wet seasons each representing 1.3% of MLRA 41-3 at SRER. For comparison, the 100 permanent field transects (with length of 30.48 m and width of 0.3 m) observes a total of 0.09 ha which is only 0.00006% of MLRA 41-3 at SRER.

The economies of scale provided by high volume drone imagery could be an appealing dataset to supplement field data collected for national-scale monitoring programs such as BLM AIM and NRI (Gillan 2020). Though it has limited ability to distinguish grass and forb species, drone imagery can expand generalized estimates of vegetation cover, provide a more robust measure of vegetation heights, and enable the development of landscape metrics not measurable from the ground. Additionally, drone imagery estimates of vegetation cover can be ‘upscaled’ to satellite imagery to cover vast landscapes (Elkind et al., 2019; Holifield-Collins et al., 2020).

All of the technologies described in this paper are available to most range practitioners in the US. Though there are some current barriers related to cost (drone equipment and software licenses), cyber infrastructure, and technical expertise, these barriers are dissolving. Drone technology and image processing software are advancing and becoming cheaper. HPC, though still housed primarily at universities and government agencies, is becoming more common and available to outside users (via web portals). Remote sensing specialists or data scientists should carry out our innovative workflow but the results and imagery products can easily by shared with less technical collaborators and stakeholders.

## Conclusion

We demonstrated a workflow to increase the efficiency of collecting, processing, and analyzing large volumes of drone imagery for rangeland monitoring applications. Our innovative workflow saved an estimated 111 workdays compared with a conventional approach. These cost savings make more practical a rich stream of monitoring data from which to link ecosystem traits with management actions. The technological barriers surrounding the use of drone imagery are quickly dissolving which will foster wider adoption by those who study and manage public rangelands.

## Acknowledgements

This material is based upon work supported by the U.S. Department of Agriculture, Agricultural Research Service, under Agreement No. 58-2022-5-13. This research was a contribution from the Long-Term Agroecosystem Research (LTAR) network. LTAR is supported by the U.S. Department of Agriculture. Any opinions, findings, conclusion, or recommendations expressed in this publication are those of the author(s) and do not necessarily reflect the view of the U.S. Department of Agriculture.

Mention of a proprietary product does not constitute a guarantee or warranty of the products by the U.S. Government or the authors and does not imply its approval to the exclusion of other products that may be suitable.

The authors declare no conflicts of interest.

## Literature Cited

Baena, S., Moat, J., Whaley, O., Boyd, D.S., 2017. Identifying species from the air: UAVs and the very high resolution challenge for plant conservation. PLoS One 12, e0188714. https://doi.org/10.1371/journal.pone.0188714

Booth, D., Cox, S., 2011. Art to science: Tools for greater objectivity in resource monitoring. Rangelands 33, 27–34. https://doi.org/10.2111/1551-501x-33.4.27

Breckenridge, R.P., Dakins, M., Bunting, S., Harbour, J.L., White, S., 2011. Comparison of Unmanned Aerial Vehicle Platforms for Assessing Vegetation Cover in Sagebrush Steppe Ecosystems. Rangel. Ecol. Manag. 64, 521–532. https://doi.org/10.2111/REM-D-10-00030.1

Breiman, J., Friedman, J., Olshen, R., Stone, C., 1984. Classification and regression trees. Chapman and Hall.

Cunliffe, A.M., Brazier, R.E., Anderson, K., 2016. Ultra-fine grain landscape-scale quantification of dryland vegetation structure with drone-acquired structure-from-motion photogrammetry. Remote Sens. Environ. 183, 129–143. https://doi.org/10.1016/j.rse.2016.05.019

D’Oleire-Oltmanns, S., Marzolff, I., Peter, K., Ries, J., 2012. Unmanned Aerial Vehicle (UAV) for Monitoring Soil Erosion in Morocco. Remote Sens. 4, 3390–3416. https://doi.org/10.3390/rs4113390

Elkind, K., Sankey, T.T., Munson, S.M., Aslan, C.E., 2019. Invasive buffelgrass detection using high-resolution satellite and UAV imagery on Google Earth Engine. Remote Sens. Ecol. Conserv. rse2.116. https://doi.org/10.1002/rse2.116

Eltner, A., Kaiser, A., Castillo, C., Rock, G., Neugirg, F., Abellan, A., 2015. Image-based surface reconstruction in geomorphometry - merits, limits and developments of a promising tool for geoscientists. Earth Surf. Dyn. Discuss. 3, 1445–1508. https://doi.org/10.5194/esurfd-3-1445-2015

Fernandez-Gimenez, M.E., McClaran, S.J., Ruyle, G., 2005. Arizona permittee and land management agency employee attitudes toward rangeland monitoring by permittees. Rangel. Ecol. Manag. 58, 344–351. https://doi.org/ht10.2111/1551-5028(2005)058[0344:APALMA]2.0.CO;2

Forlani, G., Dall’Asta, E., Diotri, F., Cella, U.M. di, Roncella, R., Santise, M., 2018. Quality Assessment of DSMs Produced from UAV Flights Georeferenced with On-Board RTK Positioning. Remote Sens. 10, 311. https://doi.org/10.3390/rs10020311

Gillan, J., Karl, J., Elaksher, A., Duniway, M., 2017. Fine-Resolution Repeat Topographic Surveying of Dryland Landscapes Using UAS-Based Structure-from-Motion Photogrammetry: Assessing Accuracy and Precision against Traditional Ground-Based Erosion Measurements. Remote Sens. 9, 437. https://doi.org/10.3390/rs9050437

Gillan, J.K., Karl, J.W., van Leeuwen, W.J.D., 2020. Integrating drone imagery with existing rangeland monitoring programs. Environ. Monit. Assess. 192, 269. https://doi.org/10.1007/s10661-020-8216-3

Gillan, J.K., McClaran, M.P., Swetnam, T.L., Heilman, P., 2019. Estimating Forage Utilization with Drone-Based Photogrammetric Point Clouds. Rangel. Ecol. Manag. 72, 575–585. https://doi.org/10.1016/j.rama.2019.02.009

Gorelick, N., Hancher, M., Dixon, M., Ilyushchenko, S., Thau, D., Moore, R., 2017. Google Earth Engine: Planetary-scale geospatial analysis for everyone. Remote Sens. Environ. 202, 18–27. https://doi.org/10.1016/j.rse.2017.06.031

Hardin, P., Jackson, M., Anderson, V., Johnson, R., 2007. Detecting Squarrose Knapweed (Centaurea virgata Lam. Ssp. squarrosa Gugl.) Using a Remotely Piloted Vehicle: A Utah Case Study. GIScience Remote Sens. 44, 203–219. https://doi.org/10.2747/1548-1603.44.3.203

Holifield-Collins, C., Skirvin, S., Winston, Z., Curley, D., Corrales, A., Armendariz, G., Gillan, J., Heilman, P., Metz, L., 2020. Improving a brush management assessment tool using drone technology and enhanced Landsat image processing, in: Society for Range Management Conference. Denver, CO.

Hugenholtz, C., Brown, O., Walker, J., Barchyn, T., Nesbit, P., Kucharczyk, M., Myshak, S., 2016. Spatial Accuracy of UAV-Derived Orthoimagery and Topography: Comparing Photogrammetric Models Processed with Direct Geo-Referencing and Ground Control Points. GEOMATICA 70, 21–30. https://doi.org/10.5623/cig2016-102

James, M.R., Robson, S., D’Oleire-Oltmanns, S., Niethammer, U., 2017. Optimising UAV topographic surveys processed with structure-from-motion: Ground control quality, quantity and bundle adjustment. Geomorphology 280, 51–66. https://doi.org/10.1016/j.geomorph.2016.11.021

Jensen, J.L.R., Mathews, A.J., 2016. Assessment of Image-Based Point Cloud Products to Generate a Bare Earth Surface and Estimate Canopy Heights in a Woodland Ecosystem. Remote Sens. 8. https://doi.org/10.3390/rs8010050

Laliberte, A.S., Rango, A., 2011. Image Processing and Classification Procedures for Analysis of Sub-decimeter Imagery Acquired with an Unmanned Aircraft over Arid Rangelands. GIScience Remote Sens. 48, 4–23. https://doi.org/10.2747/1548-1603.48.1.4

Louhaichi, M., Borman, M.M., Johnson, D.E., 2001. Spatially Located Platform and Aerial Photography for Documentation of Grazing Impacts on Wheat. Geocarto Int. 16, 65–70. https://doi.org/10.1080/10106040108542184

McClaran, M.P., Angell, D.L., Wissler, C., 2002. Santa rita experimental range digital database: User’s guide. USDA For. Serv. - Gen. Tech. Rep. RMRS-GTR 1-16. https://doi.org/10.2737/RMRS-GTR-100

Michez, A., Lejeune, P., Bauwens, S., Lalaina Herinaina, A.A., Blaise, Y., Muñoz, E.C., Lebeau, F., Bindelle, J., 2019. Mapping and monitoring of biomass and grazing in pasture with an unmanned aerial system. Remote Sens. 11, 1–14. https://doi.org/10.3390/rs11050473

Mulakala, J., 2019. Measurement Accuracy of the DJI Phantom 4 RTK & Photogrammetry.

Olsoy, P.J., Shipley, L.A., Rachlow, J.L., Forbey, J.S., Glenn, N.F., Burgess, M.A., Thornton, D.H., 2018. Unmanned aerial systems measure structural habitat features for wildlife across multiple scales. Methods Ecol. Evol. 9, 594–604. https://doi.org/10.1111/2041-210X.12919

Rehak, M., Mabillard, R., Skaloud, J., 2013. a Micro-Uav With the Capability of Direct Georeferencing. Int. Arch. Photogramm. Remote Sensing, Beijing, China XL, 4–6. https://doi.org/10.5194/isprsarchives-XL-1-W2-317-2013

Sanz-Ablanedo, E., Chandler, J., Rodríguez-Pérez, J., Ordóñez, C., Sanz-Ablanedo, E., Chandler, J.H., Rodríguez-Pérez, J.R., Ordóñez, C., 2018. Accuracy of Unmanned Aerial Vehicle (UAV) and SfM Photogrammetry Survey as a Function of the Number and Location of Ground Control Points Used. Remote Sens. 2018, Vol. 10, Page 1606–10, 1606. https://doi.org/10.3390/RS10101606

Sayre, N.F., Biber, E., Marchesi, G., 2013. Social and Legal Effects on Monitoring and Adaptive Management: A Case Study of National Forest Grazing Allotments, 1927-2007. Soc. Nat. Resour. 26, 86–94. https://doi.org/10.1080/08941920.2012.694579

Settlage, R., Chalker, A., Franz, E., Johnson, D., Gallo, S., Moore, E., Hudak, D., 2019. Open OnDemand: HPC for Everyone, in: Weiland, M., Juckeland, G., Alam, S., Jagode, H. (Eds.), High Performance Computing. Springer International Publishing, Cham, pp. 504–513.

Smith, M.W., Carrivick, J.L., Quincey, D.J., 2015. Structure from motion photogrammetry in physical geography. Prog. Phys. Geogr. 40, 247–275. https://doi.org/10.1177/0309133315615805

Snavely, N., Seitz, S.M., Szeliski, R., 2008. Modeling the world from Internet photo collections. Int. J. Comput. Vis. 80, 189–210. https://doi.org/10.1007/s11263-007-0107-3

Steele, C.M., Bestelmeyer, B.T., Burkett, L.M., Smith, P.L., Yanoff, S., 2012. Spatially explicit representation of state-and-transition models. Rangel. Ecol. Manag. 65, 213–222. https://doi.org/10.2111/REM-D-11-00047.1

Sundt, P., 2002. The statistical power of rangeland monitoring data. Rangel. J. 24, 16–20. https://doi.org/10.2458/azu_rangelands_v24i2_sundt

Toevs, G.R., Karl, J.W., Taylor, J.J., Spurrier, C.S., Karl, M.S., Bobo, M.R., Herrick, J.E., 2011. Consistent Indicators and Methods and a Scalable Sample Design to Meet Assessment, Inventory, and Monitoring Information Needs Across Scales. Rangelands 33, 14–20. https://doi.org/10.2111/1551-501X-33.4.14

West, N.E., 2003. Theoretical Underpinnings of Rangeland Monitoring. Arid L. Res. Manag. 17, 333–346. https://doi.org/10.1080/713936112

Westoby, M.J., Brasington, J., Glasser, N.F., Hambrey, M.J., Reynolds, J.M., 2012. ‘Structure-from-Motion’ photogrammetry: A low-cost, effective tool for geoscience applications. Geomorphology 179, 300–314. https://doi.org/10.1016/j.geomorph.2012.08.021

Williams, B.K., Szaro, R.C., Shapiro, C.D., 2009. Adaptive Management: the U.S. Department of the Interior Technical Guide. Adaptive Management Working Group, U.S. Department of the Interior, Washington, DC.

